# Jaccard Index Network Analysis for pangenome analysis

**DOI:** 10.1101/2025.08.20.669834

**Authors:** Arancha Peñil-Celis, Santiago Redondo-Salvo, Kaitlin A Tagg, Hattie E Webb, M Pilar Garcillan-Barcia, Fernando de la Cruz

## Abstract

Jaccard Index Network Analysis (JINA) is a comprehensive workflow designed to explore bacterial genome relationships through an integrated network-based approach. This workflow combines existing tools such as Jaccard Index (1), Gephi (2) and Pangraph (3). By integrating these methodologies into a unified framework, JINA enables efficient visualization and stratification of genomic data, facilitating the identification of meaningful patterns, groups, and associations within bacterial populations. The use of JINA ensures precision in capturing genomic variation including single nucleotide polymorphisms, insertions and deletions. While JINA does not implement Gephi, BLAST, and PanGraph directly in a single software, it guides their coordinated use to analyze and interpret genomic data effectively.

## 1 Introduction

The study of bacterial evolution is inherently complex, as it involves both vertical transmission of genetic material (inheritance from parent to offspring) and horizontal gene transfer (HGT) through mechanisms such as plasmid acquisition, recombination, or gene duplication. Addressing this complexity requires comprehensive approaches that integrate core and accessory genome elements. This integrated perspective, encompassing the pangenome, allows for a better understanding of microbial dynamics, genome adaptation, pathogenicity, and population structure.

This chapter introduces JINA as a novel workflow for stratifying bacterial populations based on genomic features from the entire pangenome. JINA is particularly useful for comparing and discriminating between very similar genomes (e.g., within a clonal serotype such as *Salmonella enterica*, serovar Typhi) because it is optimized for values well over 99.9% average nucleotide identity (ANI) (4). By combining quantitative similarity measures with network-based visualizations, JINA uncovers meaningful patterns in complex genomic datasets.

JINA leverages contemporary bioinformatic tools to analyze bacterial pangenomes and reveals genomic relationships at various levels of similarity. Central to JINA is the Jaccard Index (JI), a metric commonly used to measure similarity between datasets. In genomic applications, JI quantifies the similarity of *k*-mer sets derived from genome sequences. A *k*-mer is a contiguous subsequence of length *k*, which serves as a basic unit for genomic comparison. The JI is calculated as the ratio of the number of shared *k*-mers to the total number of unique *k*-mers across two genome sets, defined as:

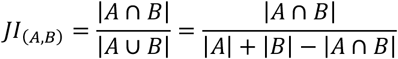

where |*A*|and |*B*| represent the number of *k*-mers in genomes *A* and *B*, respectively, and |*A*⋂*B*| represents the shared *k*-mers. JI values range from 0 to 1, with 1 indicating identical *k*-mer sets and 0 indicating no shared *k*-mers. The exact JI calculation is performed using BinDash (1), employing parameters optimized for highly similar genomes (*k*=21, minhashtype=−1) to ensure accurate results. Unlike approximations based on random subsampling, this approach considers the complete *k*-mer set, capturing subtle genomic variations such as SNPs, insertions, and deletions (indels).

JI-based network visualization and clustering is a powerful tool for pangenome exploration. It enables high-resolution stratification of thousands of genomes without requiring reference genomes, existing databases, or annotations. One significant advantage of JINA is its compatibility with both short-read and long-read sequencing data, allowing researchers to leverage the vast amount of sequencing data generated by short-read technologies.

To analyze indels more accurately, it is recommended to include at least one genome sequenced with long-read technology within each final JI-group. This strategy enhances the precise characterization of indels while reducing sequencing costs by focusing long-read efforts on representative genomes, as explained in Note 1.

The JINA workflow begins with a collection of assembled genomes, followed by calculation of the JI and the construction of an adjacency matrix to represent pairwise genome similarities. The next step involves visualizing the adjacency matrix as an undirected network. To identify groups of genomes with high sequence similarity, an optimized JI threshold is applied to filter the network. A clustering algorithm is then used to detect these groups, referred to as “JI-groups”. The final threshold-filtered network can then be stamped with relevant metadata. Lastly, the differences between JI-groups are further examined to detect indels (Figure 1).

**Figure 1:**
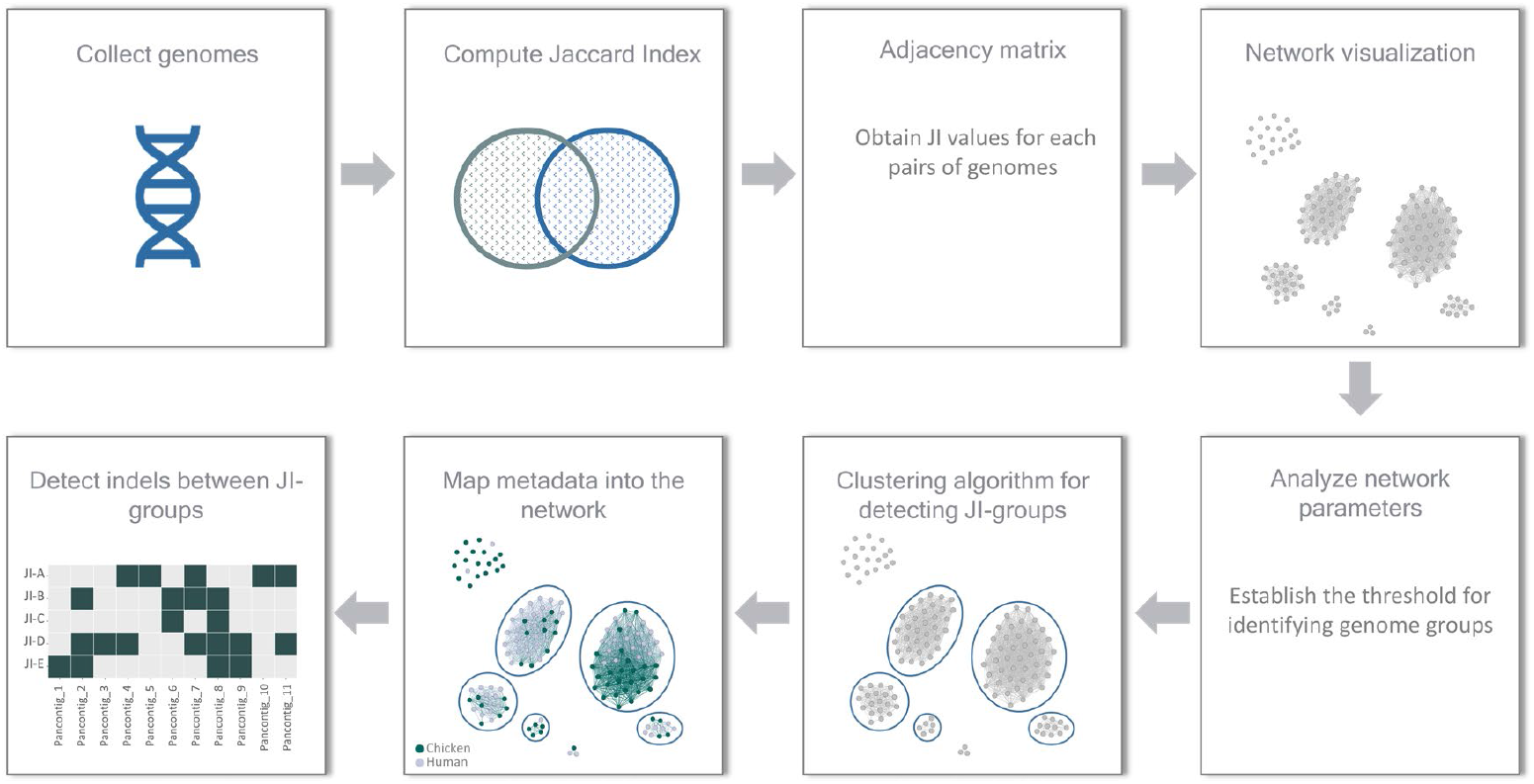
Overview of the JINA workflow. The scheme summarizes the key steps of the JINA workflow, from genome collection and pairwise similarity computation to network visualization, threshold filtering, clustering, metadata stamping, and subsequent analysis of genomic differences.

## 2 Materials

To implement the JINA workflow, the following materials, tools, and software are required:

1. **Input data**
  1.1. **Genome assemblies:** Input sequences should be assembled genomes, either from short-read or long-read sequencing technologies. It is recommended to include all genomes to be analyzed in a single folder for ease of access.
  1.2. **Genome IDs file:** Create a plain text file (genome_list.txt) containing the IDs of all genomes to be analyzed. Each genome ID should be listed on a separate line.
2. **Software and tools**
  2.1. **JINA Pipeline**. The JINA pipeline consists of a set of scripts written in Bash, Python, and Perl. Follow these steps to install:
    2.1.1. Clone the repository and adjust “PROJECT_ROOT_DIRECTORY” to your preferred location

~~~
*git clone* https://github.com/PenilCelis/MiMB_JINA_Chapter
*cd* ${*PROJECT_ROOT_DIRECTORY*}
~~~
    2.1.2. Set up the conda environment for dependencies:

~~~
*conda env create -n JINA -f JINA*.*environment*.*yml*
~~~
  2.2. **Gephi:** Install the Gephi software for network visualization. Download from https://gephi.org/
  2.3. **PanGraph:** Install the PanGraph tool for analyzing genomic differences. Follow instructions at https://github.com/neherlab/pangraph.
  2.4. **Python:** Ensure Python is installed (version 3.8 or later is recommended).
  2.5. **R:** Install R for statistical analysis. Install igraph package (https://r.igraph.org/).

## 3 Methods

This section outlines the steps required to implement the JINA workflow. Each procedure is subdivided into major steps and presented in numerical order for clarity.

1. **Genome collection:**
  1.1. Collect all assembled genomes sequences and store all genome files in a single directory for easier access during analysis.
  1.2. Generate a plain text file listing the IDs of all genomes to be analyzed, with one ID per line.
  1.3. If metadata is available, prepare it in a file where each row corresponds to a genome ID, and each column to a metadata item. Ensure the genome IDs in the plain text file and metadata file match exactly.
2. **Run JI:** This step involves converting genome assemblies into *k*-mers and calculating JI distances between each pair of genomes. The output will be an adjacency matrix represented as a list of pairwise JI values.
  2.1. **Implementation details** This process is executed using a Bash script (JINA_pipeline.sh) and the required libraries are Jellyfish (for generating and managing *k*-mer data) and BinDash (for *k*-mer similarity calculation). The input is the set of genomes and the output is the adjacency matrix containing the JI value for each pair of genomes.
  2.2. **Command to execute**

~~~
*bash JINA_pipeline*.*sh ASMB_LST ASMB_DIR OUT_DIR THREADS BLOCK_SIZE*
~~~ Replace the placeholders with the appropriate inputs for your dataset:
    - ASMB_LST: a text file containing the names of the genomes to process (e.g., “genome_list.txt”)
    - ASMB_DIR: The directory containing genome assemblies in fasta format.
    - OUT_DIR: The directory where pipeline outputs will be saved.
    - THREADS: The number of threads to use for parallel processing.
    - BLOCK_SIZE: The maximum number of genomes to compare at any time, depending on memory availability.
3. **Network visualization:** This step involves visualizing genomic relationships as an undirected network, which is constructed using the adjacency matrix obtained in step 2. Gephi v10 (https://gephi.org/) is used for network visualization, applying the ForceAtlas2 algorithm for layout. Network graphic visualization enables the exploration of clusters and relationships based on genetic similarities and genome size differences. Pairwise genome similarities can be represented in an undirected network, where nodes (genomes) are connected if the pairwise JI equals or exceeds the specified JI threshold. At the initial network stage, genomes sharing any JI value greater than 0 will be linked by an edge, resulting in most genomes forming a single connected component. By increasing the stringency of the JI threshold, separate connected components emerge (Figure 2).
  3.1. **Load data into gephi:**
    3.1.1. Import the edges.csv file generated in step 2 into Gephi as the edge list.
    3.1.2. Load metadata (if appropriate) to map node properties (e.g., geographic origin, source origin, or other genetic features).
  3.2. **Visualize the network:**
    3.2.1. Construct the network: Represent nodes (genomes) and edges (pairwise JI values). This creates an undirected network where genomes are connected if their JI meets or exceeds a given threshold.
    3.2.2. Apply the ForceAtlas2 algorithm to generate a layout that optimizes cluster visualization by positioning nodes dynamically based on their relationships. At this stage all edge connections between the nodes will be visible.
  3.3. **Filter by JI threshold:**
    3.3.1. Start with an inclusive network (JI > 0). This will create a highly connected network where most genomes are part of a single component (first panel of Figure 2).
    3.3.2. Gradually increase the JI threshold to leave only edges between genomes with high similarity. At each step, reapply the ForceAtlas2 algorithm. This allows the nodes to dynamically reposition themselves, adapting to the evolving structure of the network and highlighting changes in relationships as the threshold becomes more stringent. Observe the network as tighter clusters emerge, corresponding to genomes that are more closely related. Continue this iterative process (adjusting the threshold) until the network exhibits distinct, meaningful clusters: highly internally connected subgraphs (groups of genomes with strong internal similarity), reduced connectivity between separate groups and, strive to retain a majority of genomes in well-defined, non-singleton communities to provide informative insights.

**Figure 2:**
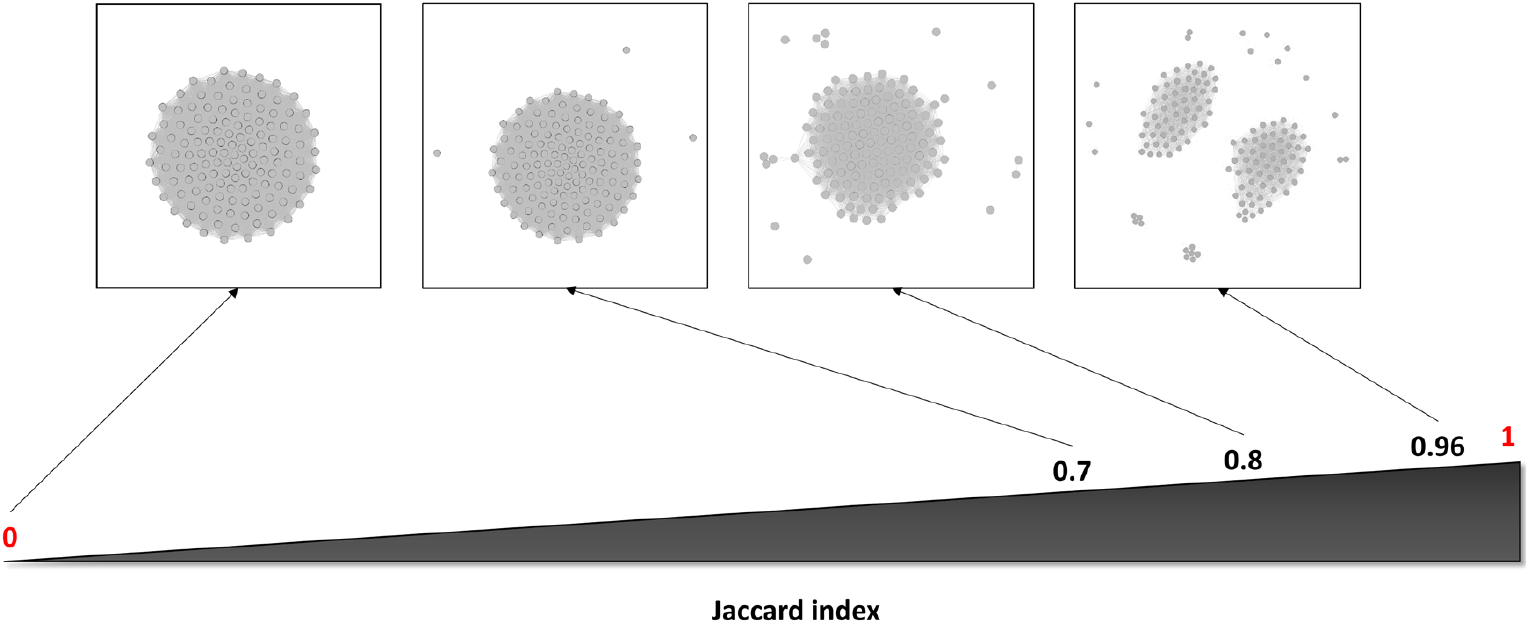
Example of a network representation based on pairwise genome similarities. Nodes represent genomes, and edges are drawn when the pairwise JI value meets or exceeds the defined threshold. The initial network includes all genomes connected by any JI value above 0, forming a single component, while increasing the JI threshold (0.7, 0.8 and 0.96) results in the emergence of separate connected components. The represented network corresponds to a group of 135 genomes of *Salmonella enterica* serovar Enteritidis. The most convenient JI threshold depends on the specific goals of your project and the dataset being analyzed. Refer to Step 4 (“Analyze network parameters”) for additional guidance in choosing the most suitable threshold. See also Note 2 for further details.
4. **Analyze network parameters:** The analysis of network parameters is helpful for determining the most appropriate JI threshold for identifying genomic groups in your dataset. Metrics such as transitivity and density provide insights on network’s structure and strength of its communities.
  4.1. **Implementation details:** The analysis can be performed in R using the igraph package. The required input is the edges.csv file, which contains the pairwise adjacency matrix of JI values. Functions from igraph allow the calculation of key network metrics that guide the threshold selection process.
  4.2. **Threshold optimization:** The choice of the JI threshold determines the level of similarity required for genomes to be grouped, influencing the sparsification of the network and the emergence of distinct communities.
    4.2.1. The most appropriate threshold depends on the specific goals of the project and the dataset being analyzed. For instance, if the goal is to differentiate highly similar genomes, such as those belonging to the same serovar of *Salmonella enterica*, you may want to use a high JI, well above 0.98 (4). Conversely, for more diverse datasets or broader genomic comparisons, a lower JI threshold may be more suitable to capture meaningful but less stringent similarities.
    4.2.2. Several statistical parameters can be evaluated to guide the optimization process:
      - Transitivity: Indicates the strength of internal connectivity within communities. High transitivity indicates that the nodes within communities are highly connected, forming tightly-knit groups. While high transitivity suggests strong communities, it may not always align with the optimal JI threshold because a threshold based on very high transitivity might result in too many small clusters, which could be impractical for downstream analysis.
      - Number of communities: Balance the number of clusters to ensure biological relevance. Over-fragmented networks with many small communities may not be informative. As a guideline, aim for a number of clusters that reflects the dataset’s diversity without exceeding practical limits for analysis. As a rule of thumb, keep the total number of groups to less than the square root of *N, N* being the total number of nodes in the network. Communities with fewer than five members are generally less informative.
      - Proportion of genomes in communities: Ensure that most genomes belong to non-singleton communities. Singletons or poorly connected nodes reduce the interpretability of the network.
      - Incorporate knowledge of what specific JI thresholds mean in terms of genomic differences, as explained in Peñil-Celis et al., 2024 (4).
      - Reflect congruence with genetic determinants or metadata of interest, where available. Congruence between the resulting clusters and known genetic determinants or metadata (e.g., geographic origin, serovar, or AMR profiles) can provide valuable guidance in selecting the most appropriate JI threshold.
5. **Cluster detection using the Louvain algorithm**. The purpose of this step is to classify the genomes into JI-groups. We propose two options:
  5.1. **Gephi plugin**. Gephi provides a Louvain clustering algorithm as a plugin for community detection. Steps in Gephi:
    5.1.1. Apply the Louvain algorithm from the “Statistics” tab.
    5.1.2. Analyze the resulting communities to identify JI-groups.
  5.2. **Igraph package in R:** Alternatively, use the igraph library in R (https://igraph.org) to perform Louvain clustering. Steps:
    5.2.1. Load the network data (edges.csv) into R and create an igraph object (basic_net).
    5.2.2. Use the cluster_louvain() function to perform clustering.
    5.2.3. Extract and analyze the resulting clusters to define JI-groups. See Note 3 for further guidance on Louvain algorithm.
6. **Stamp metadata into the network:** The network nodes, representing genomes, can be stamped (e.g., colored) based on metadata and genetic determinants of interest. Metadata stamping allows for visual evaluation of associations between the identified groups and various epidemiological data, genetic factors, or other relevant variables. By integrating these layers of information, network visualization might reveal potential correlations and patterns, highlighting relationships that may guide further analyses. Moreover, metadata can assist in selecting the optimal JI threshold.
7. **Detect indels between JI-groups**. The identification of indels, including MGEs and other accessory genome regions, between JI-groups is proposed to be carried out using PanGraph (3). PanGraph is specifically designed to analyze the pangenome. It provides a command-line interface to identify homology among large collections of closely related genomes. PanGraph identifies blocks of homologous sequence and it is used to detect indels specific to each JI-group.
  7.1.1. **Prepare input:** The input for PanGraph must be a single FASTA file where the header corresponds to the ID of each genome. Concatenate all genome assemblies into a single compressed file (e.g.,.fa.gz). This concatenated file will be the input of PanGraph.
  7.1.2. **Run PanGraph (Figure 3A):** Execute PanGraph on all genomes using the appropriate parameters as detailed in the official documentation. Basic usage would be: *pangraph build sequence*.*fa*.*gz > graph*.*json*, where sequence.fa.gz is a multifasta with all genomes to analyze and graph.json is the output graph file representing the pan-genome structure. Then, export the graph to obtain data for further analysis: *pangraph export graph*.*json*
  7.1.3. **Process PanGraph output using PyPanGraph (Figure 3B)**. Use the PyPanGraph Python library (https://github.com/mmolari/pypangraph/) to process the output from PanGraph. This library was specifically designed to provide utilities for interacting with pan-genome graphs and extracting useful data from PanGraph output. Key outputs of this library include: CSV File with pancontig data that contains details such as the length of each pancontig, the number of genomes in which a pancontig appears, and the presence/absence matrix, showing the presence or absence of each pancontig in the analyzed genomes.

**Figure 3:**
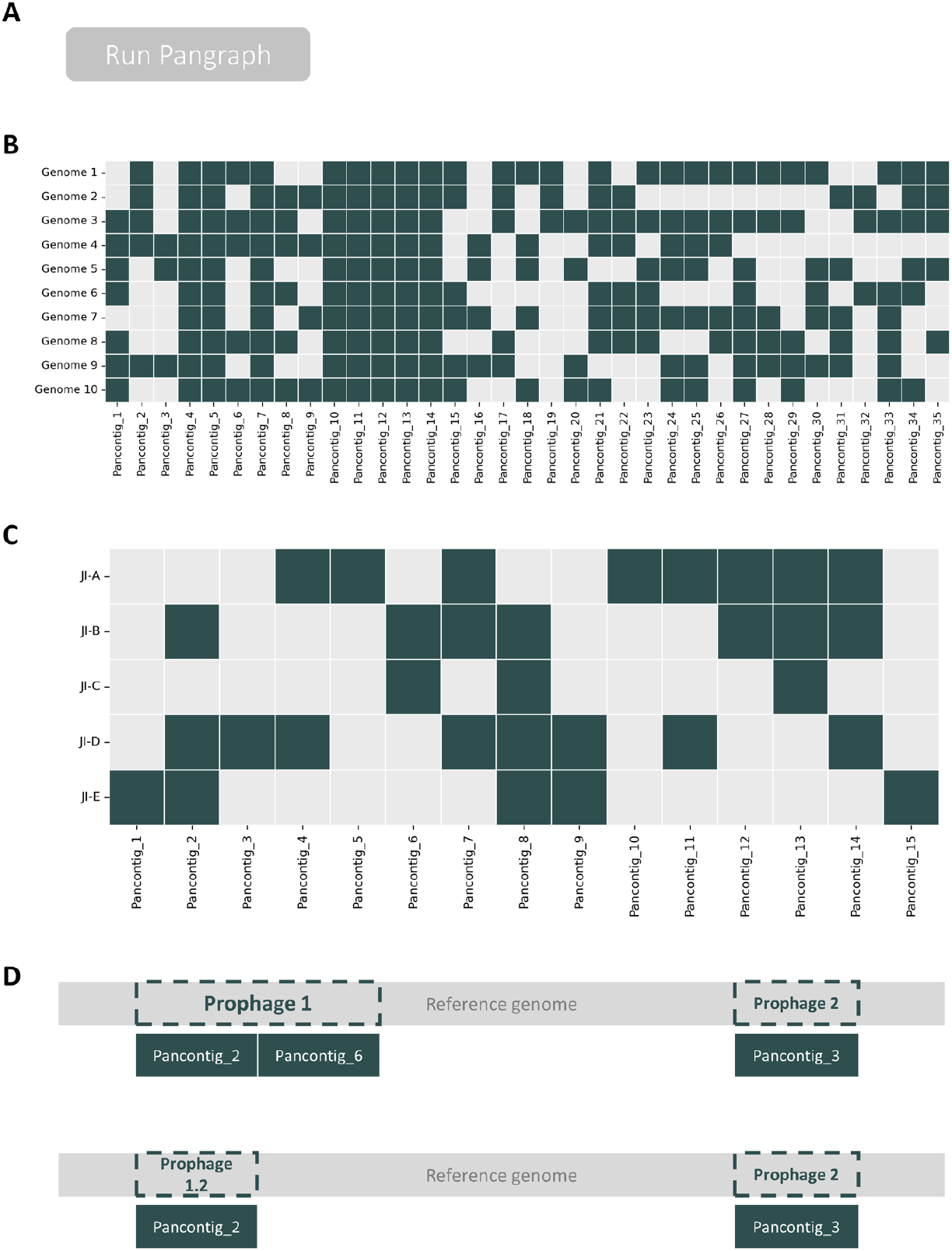
Detection of JI-group specific indels using PanGraph. (A) **Run PanGraph**. (B) **Presence/Absence Matrix**: A matrix is generated to display the presence (green cells) or absence (gray cells) of pancontigs across all genomes. (C) **Classification into JI-groups:** Pancontigs that meet the threshold for defining “core” pancontigs in each JI-group (for instance 90% of the members of each JI-group and absent from other JI-groups) are retained for further analysis. This process generates a matrix with all JI-groups, where the core pancontigs of each JI-group are indicated (green cells). (D) **Stamping core pancontigs to a reference genome:** The core pancontigs are stamped with BLASTn against a reference genome to arrange them and detect the regions they form. For example, pancontig_2 and pancontig_6 are found to be consecutive, and they encode phage-related proteins so they are considered to form a unique prophage.
  7.1.4. **Identify core pancontigs specific to each JI-group (Figure 3C)**. Start with the presence/absence matrix generated in 7.1.3 and the classification of genomes into JI-groups and divide the matrix into separate matrices for each JI-group. Within each JI-group matrix, filter the pancontigs based on criteria that best suit your analysis: for example, you might remove small pancontigs (such as those under 250 bp) to focus on longer, more informative sequences, and adjust the threshold for defining characteristic pancontigs (e.g., requiring their presence in at least 90% of the genomes in each JI-group). Exclude any pancontigs present in all genomes across all JI-groups, as they do not provide group-specific information. After filtering, compile the core pancontigs from all JI-group into a single output, such as a csv file or a heatmap (Figure 3C). In this output the first column lists JI-groups and the subsequent columns correspond to the core pancontigs. Each cell is filled to indicate the presence or absence of a given pancontig in a specific JI-group.
  7.1.5. **Map core pancontigs to a reference genome (Figure 3D):** Use BLAST to map core pancontigs against a reference genome (preferably sequenced using long-read technology, see Note 1) from the same JI-group. This enables correct ordering of pancontigs and facilitates the reconstruction of their original arrangement within the genome. This step helps identify genomic regions composed of multiple consecutive pancontigs, such as a prophage composed of several adjacent pancontigs, thereby allowing for accurate reconstruction of such elements

## 4 Notes

1. Importance of reference genomes for accurate pancontig analysis. In JINA, the use of reference genomes sequenced with long-read technology (or retrieved from public archives such as National Center for Biotechnology Information (NCBI, https://www.ncbi.nlm.nih.gov/), is strongly recommended for accurate mapping and characterization of core pancontigs. Selecting at least one genome sequenced with long-read technology for each JI-group ensures that representative genomic features are captured while minimizing sequencing costs. This is particularly important because accessory genome elements are often composed of multiple consecutive pancontigs (3). Long-read sequencing provides the continuity needed to identify and correctly order these consecutive pancontigs, ensuring accurate reconstruction of such regions.
2. Interpret JI values. For detailed insights into what specific JI values represent in terms of insertions, deletions (indels), or SNPs, refer to the publication by Peñil-Celis et al., 2024 (4), which provides a comprehensive breakdown of the implications of different JI thresholds. For example, a JI threshold of 0.983 can group genomes that differ in indels up to 100 kb in size, 2200 SNPs, or a combination of both.
3. Louvain clustering. The Louvain algorithm optimizes modularity to detect communities with dense internal connections while minimizing links between groups. The resolution parameter is key to controlling the granularity of the clusters.
  3.1. In R (igraph Implementation):
    - Higher resolution: Results in more, smaller clusters, making it useful for detecting highly similar genomes.
    - Lower resolution: Produces fewer, larger clusters, which is better suited for broader comparisons.
  3.2. In Gephi: The behavior of the resolution parameter is inverted compared to R.
    - Higher resolution in Gephi: Produces fewer, larger clusters.
    - Lower resolution in Gephi: Results in more, smaller clusters.

## 5 Disclaimer

The findings and conclusions of this report are those of the authors and do not necessarily represent the official position of the Centers for Disease Control (CDC).

